# Utilization and degradation of laminarin-based substrates by marine yeasts suggests their niche-specific role in microbial loop dynamics

**DOI:** 10.1101/2024.09.13.612705

**Authors:** Berin S. Arslan-Gatz, Mikkel Schultz-Johansen, Tom-Niklas Hollwedel, Sofie Niggemeier, Rolf Nimzyk, Antje Wichels, Gunnar Gerdts, Jan-Hendrik Hehemann, Tilmann Harder, Marlis Reich

## Abstract

In the oceans, the diversity of phytoplankton primary products supports a wide range of microbial heterotrophs, including bacteria and fungi. The organic substrate dynamics within pelagic microbial communities are strongly controlled by microorganismal interactions, resulting in a dense interactome. While the role of bacteria in the microbial loop is well documented, the degradation capacity and substrate specificity of marine fungi, as well as their role and function in metabolic guilds with bacteria, is comparatively less understood. We chose the polysaccharide laminarin, a major product of marine primary production, as well as oligomeric laminarin subunits and monomeric glucose, to study the degradation capacity of eleven marine yeast isolates from the pelagic microbial community of Helgoland Roads. Our aim was to measure yeast growth and correlate degradation yields and putative intermediate degradation products with the size of laminarin-based organic precursor substrates. We developed a reproducible, temporally resolved, high-throughput growth protocol to measure resource-specific yeast growth. Measurement of temporally fine-scaled growth kinetic models of isolates were accompanied with qualitative and quantitative chemical analyses of substrates and degradation intermediates. Our data showed that yeast growth was negatively correlated with oligomer length. Fluorophore-assisted carbohydrate electrophoresis suggested the lack of enzymatic endo-activity for laminarin in yeasts under investigation, suggesting they may occupy a niche in the microbial loop, benefitting from extracellular hydrolysis of carbohydrates by other microorganisms. In terrestrial environments, namely forest soil ecosystems, yeasts have been assigned a similar niche, supporting a prominent role of yeasts in microbial interactomes.

## 1 Introduction

Marine phytoplankton produce a complex pool of organic material, especially during bloom scenarios [1]. The majority of this material is immediately utilised and transformed at the base of the food web by heterotrophic microbial consortia, such as bacteria and microeukaryotes [2]. This process, known as the "microbial loop" [3], results in the assimilation and respiration of organic matter otherwise inaccessible to higher trophic levels of the marine food chain.

With their high chemical diversity and different stages of solution, adsorption and aggregation, marine organic compounds differentially support a wide range of microbial heterotrophs. The temporal and qualitative dynamics of phytoplankton bloom-derived organic material [4] thus affect pelagic microbial community dynamics. Next to substrate quality and quantity, these dynamics are largely structured by organismal interaction during and post phytoplankton blooms, e.g., by competition or resource sharing among community members [5] resulting in a dense microbial interactome [6, 7].

The bacterial contribution to the microbial loop is well studied, and reproducible succession patterns of specific bacterial clades have been observed and explained by substrate-induced forcing during and after phytoplankton blooms [4]. The bacterial decomposition strategies, species-specific enzyme repertoires [8] and feeding strategies [9] strongly control the role of individual bacteria within the interactome. In contrast, marine heterotrophic fungi have long been neglected as potential members and potential drivers of the microbial loop, despite their ubiquitous distribution, functional diversity and abundance [10, 11]. Based on few studies, seasonal dynamics of marine fungi correlate with nutrients and organic matter as well as the presence of interaction partners, e.g., phytoplankton, zooplankton and bacteria [12–14]. Notably, during phytoplankton blooms, both free-living saprotrophic filamentous fungi [15] and yeasts [16] reach ecologically relevant biomass, supporting the notion of fungi as relevant players in the microbial loop. The identification of specific carbohydrate-active fungal enzymes (CAZymes) [17] with a high functional diversity in global pelagic omic datasets as well as stable-isotope probing (SIP) analyses of pelagic communities [18, 19] further support their active involvement in the microbial loop.

However, compared to bacteria, controlled and time-resolved *in vitro* cultivation studies of marine fungi under growth regimes and media compositions mimicking pelagic environments are largely lacking (but see [11, 20–22]), especially for relevant fungal members in natural microbial communities. Yet, such experiments are urgently sought to differentiate substrate utilization characteristics and kinetics among fungi through growth and chemical analyses of substrate degradation, and unravel their role and importance in the microbial loop. Thus, one goal of this study was to establish reproducible time-resolved high-throughput protocols to detect substrate-specific fungal growth.

The organic matter released during phytoplankton blooms comprise a broad spectrum of chemical compound classes. High-molecular-weight (HMW) components, such as polysaccharides, make up a large proportion of dissolved organic carbon (DOC) [23]. One major DOC constituent is the helical polysaccharide laminarin, which consists of glucose monosaccharides forming a β-1,3-linked backbone and β-1,6-linked branches. Since HMW laminarin is a ubiquitous internal energy storage compound in photosynthetic eukaryotes it is widely abundant in phytoplankton blooms, accounting for up to a quarter of the annual marine primary production of 49 gigatons of carbon, and 3-digit micromolar concentrations in ambient seawater [24, 25].

The second goal of this study was to use the optimal growth protocol to analyse the relationship of organic substrate size on fungal degradation kinetics (Figure 1). In accordance with previous marine bacterial carbohydrate degradation studies [26], we chose HMW laminarin as ecologically relevant model organic substrate. Next to HMW laminarin we included enzymatically partially degraded laminarin, oligomeric laminarin subunits and monomeric glucose in degradation studies. Specifically, we analysed the temporally resolved degradation capacity of these substrates with up to eleven yeast species, all of which occur and have been isolated in the pelagic microbial community at the time series station of Helgoland Roads, North Sea, Germany [14, 16].

**Figure 1:**
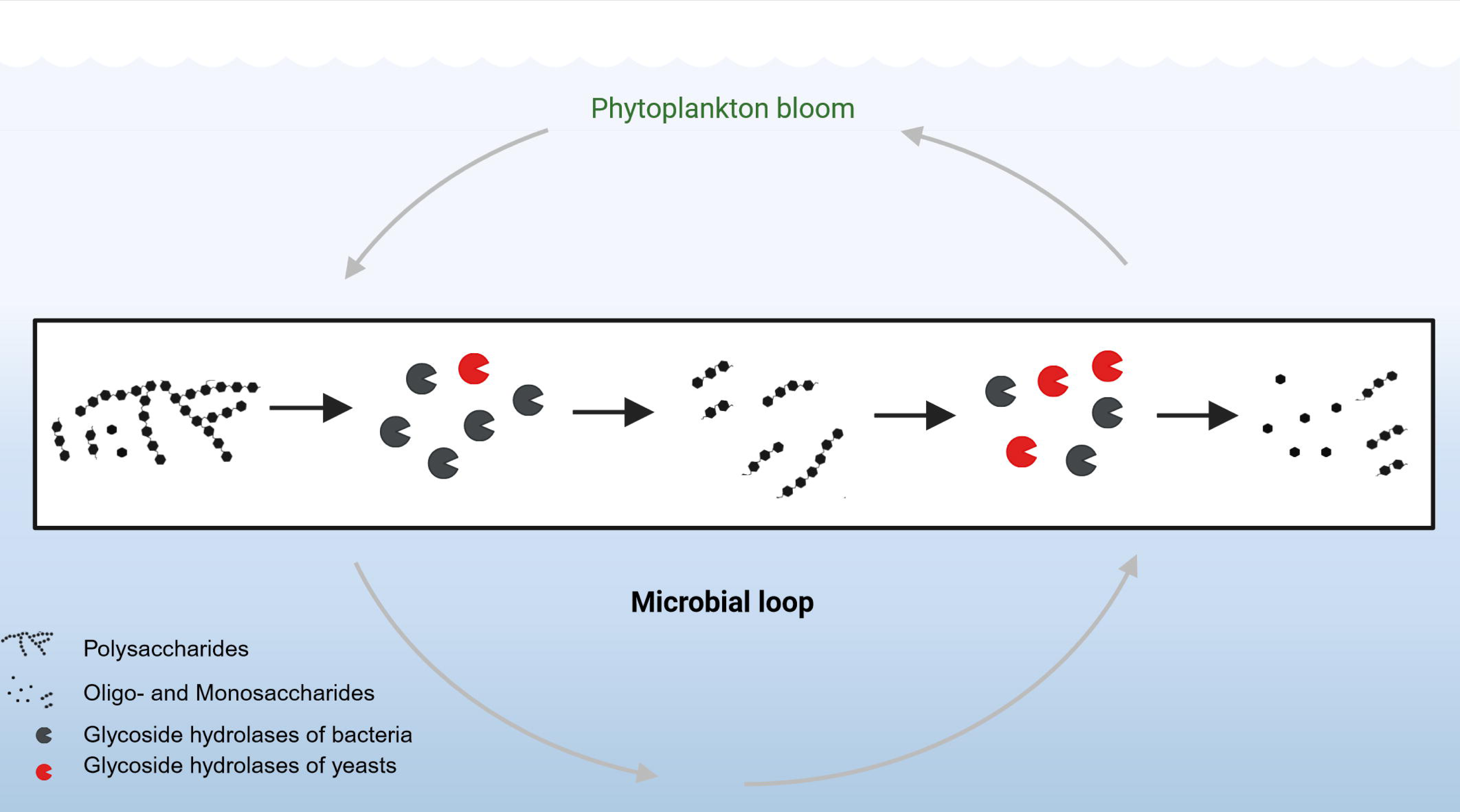
Graphical abstract: Niche position of marine yeasts in the microbial loop: This graphical abstract depicts the proposed role of marine yeasts within the marine microbial loop, highlighting their interactions with bacteria and phytoplankton. The box indicates the flow of organic matter and energy, with marine yeasts contributing to nutrient cycling and energy transfer by largely degrading oligosaccharides, thereby facilitating nutrient availability. Created in BioRender. Niggemeier, S. (2024) BioRender.com/o73b845. [56].

## 2 Materials and Methods

### 2.1 Fungal isolates

The eleven marine yeast isolates used in this study stemmed from the fungal strain collections of M. Reich (unpublished), A. Wichels and G. Gerdts [27] (Table 1). All isolates originated from surface water (1 m) of the Long-Term Ecological Research (LTER) station Helgoland Roads (54°11.3’ N, 7°54.0’ E). The selection of isolates was based on taxonomy, abundance and occurrence in the mycoplankton community of Helgoland Roads, as detected during the spring phytoplankton bloom in 2017 [16] (Figure 2) and over the course of 2015/2016 [14] (Table 1).

**Figure 2:**
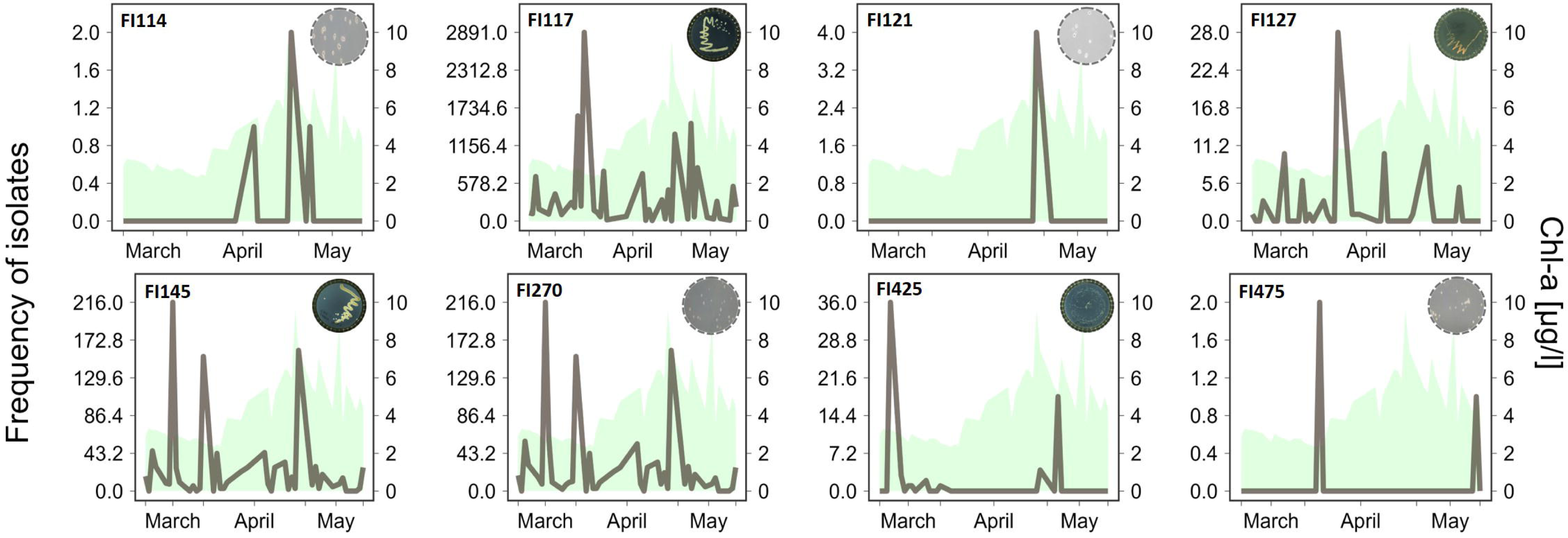
Occurrence of yeast isolates in surface water from Helgoland Roads during the spring phytoplankton bloom of 2017 [ref. 16]. The frequency of the individual isolates (based on OTU sequence frequency) is shown in relation to chlorophyll a content in water, which is used as a proxy for phytoplankton cell numbers. Isolate code in the upper left corner of each graph.

**Table 1:** Taxonomic information and occurrence of eleven yeast isolates in environmental datasets. Isolation source of all isolates was the surface water (1m) of Helgoland. A, Ascomycota; B, Basidiomycota.

| Isolate ID | Taxonomy | Species hypothesis (UNITE<sup>1</sup>) | Taxonomy | Category of occurrence pattern in environment as Banos et al., 2020<sup>2</sup> | Present in spring phytoplankton bloom as Priest et al., 2021<sup>3</sup> | Reference |
| --- | --- | --- | --- | --- | --- | --- |
| FI018 | *Metschnikowia bicuspidata* | NA | Saccharomycetes (A) | IV - long-lasting | NA | culture collection Reich lab |
| FI113 | *Cystofilobasidium infirmominiatum* | SH1295646.09FU | Tremellomycetes (B) | NA | NA | Krause et al., 2013<sup>4</sup> |
| FI114 | *Leucosporidium scottii* | SH4274207.09FU | Microbotryomycetes (B) | IV - long-lasting | yes | Krause et al., 2013<sup>4</sup> |
| FI117 | *Protomyces* sp. | SH1022425.09FU | Taphrinomycetes (A) | III - steady | yes | Krause et al., 2013<sup>4</sup> |
| FI121 | *Vishniacozyma victoriae* | SH0994206.09FU | Tremellomycetes (B) | IV - long lasting | yes | Krause et al., 2013<sup>4</sup> |
| FI122 | *Hyaloscyphaceae* sp. | NA | Leotiomycetes (A) | II - frequent peaking | NA | Krause et al., 2013<sup>4</sup> |
| FI127 | *Rhodotorula mucilaginosa* | SH1008722.09FU | Microbotryomycetes (B) | IV - long-lasting | yes | culture collection Reich lab |
| FI145 | *Candida* sp. | NA | Saccharomycetes (A) | II - frequent peaking | yes | culture collection Reich lab |
| FI270 | *Candida sake* | SH1040972.09FU | Saccharomycetes (A) | I - boom-bust like | yes | culture collection Reich lab |
| FI425 | *Kondoa aeria* | SH1116283.09FU | Agaricostilbomycetes (B) | III - steady | yes | culture collection Reich lab |
| FI475 | *Tausonia pullulans* | SH0753078.10FU | Tremellomycetes (B) | II - frequent peaking | yes | Krause et al., 2013<sup>4</sup> |

<sup>1</sup>Köljalg et al., 2020; DOI: 10.3390/microorganisms8121910
<sup>2</sup>Banos et al., 2020, DOI:10.3389/fmicb.2020.01305
<sup>3</sup>Priest et al., 2021; DOI:10.1111/1462-2920.15331
<sup>4</sup>Krause et al., 2013; DOI:10.1007/s10152-013-0348-1
A = Ascomycota; B = Basidiomycota

### 2.2 Precultures of yeast isolates

For each isolate, new precultures were prepared from glycerol cryo stocks using artificial seawater (ASW) [28, 29] and YM medium [30] containing 100 mg/ml each of ampicillin and kanamycin (Roth, Karlsruhe, Germany). Subsequently, the medium was inoculated with the cryo stock and cultivated at 16°C under a 13/11 h day/night cycle at 150 rpm on a KL 2 shaker (Edmund Bühler GmbH, Tübingen, Germany) until visible growth. Thereafter, cultures were inoculated at 1:1 ratio into fresh medium to a final volume of 1.5 ml. After three days, 750 µl culture were transferred into 1:4 diluted YM medium to a total volume of 20 ml. After another three days, these cultures served as inoculum for growth studies described below.

### 2.3 Growth experiments

Growth experiments were conducted in 96-well plates (CELLSTAR, Greiner Bio-One GmbH, Frickenhausen, Germany) in 200 µl volume per well at 16°C and 260 rpm (CMS1000, Cerillo, Charlottesville, VA, USA). The initial optical density (OD) was set to 0.1. Each isolate was grown in replication (n = 4). A total of 12 wells per plate were used as ASW blanks. The growth of yeasts on internal storage products only was measured in ASW without further additions (n = 4). Growth was monitored by automated OD recordings at 600 nm over 65 h at 30 min intervals using an ALTO plate reader (Cerillo, Charlottesville, VA, USA).

The culture protocols were developed on three yeast isolates FI018, FI114 and FI117 with the exception of growth conditions (B), (D), and (F) (see below), for which FI018 was replaced with FI113. Preliminary growth experiments were carried out with glucose (Roth, Karlsruhe, Germany) as sole carbon source at 2.5 g/l. The other conditions were adapted incrementally to the following parameters:

A. Different concentrations of inorganic nitrogen (N) and phosphorous (P) required for yeast growth were tested. Naturally occurring N and P concentrations at Helgoland Roads during the 2016 spring phytoplankton bloom [31] served as a reference and were tested at 10-fold and 100-fold elevated concentrations. These settings were trialled with 75 µM and 30 µM, and 750 µM and 300 µM of NaNO_3_ and NaH_2_PO_4_, respectively.
B. As the potential utilisation of organic carbon by yeasts is governed by the stoichiometric ratio of C:N:P [32], we tested growth at different stoichiometric C:N:P ratios of growth media. The P concentration of 30 µM determined in (A) served as the baseline. Elements were supplied in C:N:P ratios of 106:16:1 (Redfield ratio) and higher (173:16:1) and lower (17.3:16:1) proportions of carbon adjusted by 2.5 g/l and 0.25 g/l glucose. Growth was compared to control media containing the same glucose concentrations, but with N and P concentrations of 75 µM and 30 µM, respectively, as in (A), resulting in a stoichiometric ratio of 17.3:2.5:1 and 173:2.5:1.
C. As the environmental pH value may affect the efficiency of extracellular enzymes, such as laminarinase [33], we tested fungal growth in ASW at pH 7.0 and enriched ASW (EASW) [28, 29] at pH 8.2.
D. To distinguish between fungal growth on internal energy storage products and organic resources added to growth media, a nutrient deprivation step was introduced prior to growth experiments. Yeast cells from precultures at OD 0.6 to 1.2 were transferred into ASW to a final OD of 0.4. Nutrient deprivation of 45 and 65 hours were compared to growth without deprivation.
E. To assess growth detection thresholds in media containing glucose or HMW laminarin as sole carbon source, growth was measured in a 10-fold glucose concentration series from 0.00025 to 25 g/l; growth was still observed at 0.025 g/l glucose. The detection threshold of HMW laminarin (ca. 7,500 Da, 10 repeating units) was tested at 0.017, 0.17, and 0.97 g/l, corresponding to 0.025, 0.25 and 0.85 g/l glucose equivalent as determined by the phenol-sulfuric acid (PSA, see section 2.4) quantification.
F. Since expression of enzymatic degradation pathways consumes initial energy, especially in the case of complex and structurally rich polysaccharides, we distinguished growth of the same yeast isolates under “primed” conditions, i.e., the isolate was raised on the carbon source under investigation prior to growth studies, and “non-primed” conditions, i.e., the isolate was exposed to the substrate at the beginning of the experiment. To prime a preculture, 0.0017 g/l HMW laminarin was added during the nutrient deprivation period. Growth experiments with yeast isolates obtained under different pre-treatment conditions were carried out at the lowest HMW laminarin concentration at which growth was still detected for all isolates (as tested in (E)).

### 2.4 Size definition of HMW laminarin and total carbohydrate quantification in laminarin-based media

A stock solution of 1.8 g/l laminarin was prepared (>96% purity, from *Laminaria digitata,* L9634, Sigma-Aldrich®, Germany) and sequentially size-fractionated with spin column filters (Sartorius, Göttingen, Germany) of 5 and 10 kDa MWCO. The working solution used in all experiments contained laminarin in the size range of 5 to 10 kDa, in the following referred to as HMW laminarin. The carbohydrate concentration in HMW laminarin-based media was determined with the phenol-sulphuric acid (PSA) method [34] with modifications. A HMW laminarin calibration curve was established in the range of 0.02 - 0.9 g/l (Suppl. Fig. 1). Briefly, 550 µl of HMW laminarin working solutions were transferred to 1.5 ml reaction tubes, topped with 367 µl 10 M hydrochloric acid, and carefully vortexed. Samples were hydrolyzed at 100 °C for 2 h under orbital shaking at 600 rpm (ThermoMixer C, Eppendorf, Hamburg, Germany). After cooling, 120 µl of hydrolyzed samples were transferred into new reaction tubes and 600 µl of 95% sulfuric acid were added. Samples were heated to 90°C for 15 min at 900 rpm followed by addition of 120 µl 5% aqueous phenol. The reaction mixture was vortexed for 2 min, stored for 1 h, after which the absorbance at 490 nm was measured in 96 well plates (200 µl per well) in a plate reader (Clariostar Plus, BMG Labtech, Germany) with technical replication (n = 3).

### 2.5 Preparation of partially hydrolyzed laminarin

Two laminarinases, FaGH17A and FbGH30 from *Formosa* spp. (marine flavobacteria, see [35]) were used to partially digest HMW laminarin prior to growth experiments. FaGH17A is an endo-acting enzyme that hydrolyses β-1,3-linkages of laminarin, while FbGH30 cleaves the β-1,6-linked glucose branches along the laminarin backbone. Both enzymes have high specificity for laminarin at pH 7.0 [33]. A phosphate buffered saline (137 mM NaCl, 2.7 mM KCl, 10 mM Na_2_HPO_4_, 1.8 mM KH_2_PO_4_) HMW laminarin stock solution (4 g/l) at pH 7.0 was hydrolysed for 16 h at RT with 0.5 µM of each of the two enzymes in a final volume of 1.6 ml. The enzymes were inactivated by incubation at 98°C for 10 min, followed by centrifugation at 16,000 x g for 10 min at 4°C. The supernatant was collected and the partially hydrolysed laminarin solution was sterile-filtered, quantified by PSA and diluted to a working concentration of 0.85 g/l.

### 2.6 Yeast growth experiments with HMW laminarin and partially hydrolyzed laminarin as single carbon source

The growth of 11 yeast isolates on HMW laminarin and partially hydrolyzed laminarin (size unknown) as only carbon source at a normalized concentration of 0.85 g/l glucose equivalents (PSA) was compared. The positive control contained glucose at a concentration of 0.85 g/l, which represents the glucose equivalent of 0.97g/l HMW laminarin (ca. 7500 Da, 10 repeating units), while the negative control contained ASW medium only.

### 2.7 Tracking and quantification of substrates and intermediate degradation products over time

To track the degradation of laminarin-based precursor substrates over time and identify intermediate degradation products, samples were taken at 5 time points (n =1) and profiled with fluorophore-assisted carbohydrate electrophoresis (FACE) or quantified by PSA (see 2.4) using three isolates (FI122, FI425 and FI475). In addition to HMW laminarin and partially hydrolyzed laminarin, a mixture of two laminarin oligosaccharides (laminarihexaose and laminaribiose; Megazyme, Auchincruive, UK) was used in the standard growth protocol (Supplementary S4).

Specifically, subsamples (25 µl) were taken at five time points, zero (T0), 9 (T1), 22 (T2), 45 (T3), and 70 h (T4). Enzyme activity in these samples was immediately quenched by boiling at 99°C for 10 minutes. Samples containing only HMW laminarin, with a total volume of 25 µl, were dried under vacuum, redissolved in 5 µl of ultrapure water, and then derivatized according to Jackson et al. [36]. For the other samples, including partially hydrolyzed laminarin and mixtures of laminarin oligosaccharides, only 5 µl of the subsample was used for the derivatization reaction. Briefly, 1 µl 0.02 M 8-aminoaphtalene-1,3,6-trisulfonic acid (ANTS) and 2 µl 1 M NaBH_3_CN were added to each 5 µl sample. The reaction mixture was incubated overnight at 37°C. Three microliters of 100% glycerol were added and the total of 8 µl was loaded onto a standard acrylamide gel (35%). Electrophoresis was performed at 100V for 30 min followed by 200V at 4°C using pre-cooled running buffer (25 mM Tris-base, 250 mM glycine). An oligosaccharide standard was prepared by combining 0.85 mg/ml laminarihexaose, 0.85 mg/ml laminaripentaose, 0.56 mg/ml laminaritetraose, 0.43 mg/ml laminaribiose (all from Megazyme), and 0.21 mg/ml glucose (Sigma).

### 2.8 Statistics

The computer program AMIGA (download 2023-12-05) [37] was used to analyze the growth data and model the growth curves. Growth curves were modelled by non-parametric Gaussian Process (GP) regression approach. Firstly, each tested factor was checked for significant differences between growth on internal storage materials (negative control) versus the carbon source provided using the Bayesian test with the default settings [38]. In case of significant differences, growth models were normalized to the negative control using the subtraction method. Based on the models, AMIGA estimated parameters describing the growth type such as the normalized maximal OD (norm ODmax).

To test differences due to growth conditions or resource-specific growth, Kruskal-Wallis test was performed on individual growth parameters and resources, followed by Dunn’s test of multiple comparisons using a p-value with Bonferroni correction of ≤ 0.05 as threshold for statistical significance. Both tests were run with the RStudio program v 2024.04.0+764 using the core functions as well as the ‘Dunn.test’ package [39]. Graphs were designed by using the package ‘ggplot2’ [40].

## 3 Results

### 3.1 Optimal growth parameters

The preliminary tests of different growth conditions resulted in the following protocol. All growth experiments were conducted in 96-well plates over a period of 70 h. ASW served as a medium base, supplemented with 75 µM NaNO_3_, and 30 µM NaH_2_PO_4_, while the C:N:P ratio was not adjusted to maintain the Redfield ratio. Precultures of the fungal isolates were subjected to a nutrient deprivation period of 45 h without any priming prior to the growth experiment with different carbon sources. To ensure best growth among all yeast isolates, 0.85 g/l glucose equivalents of HMW laminarin, partially hydrolyzed laminarin, and laminarin oligosaccharides were used as the carbon source, while same amount of glucose was used as the positive control. For details of the stepwise parametrization of this growth protocol see Supplementary S1.

### 3.2 Yeast growth on HMW laminarin, partially hydrolyzed laminarin and glucose

All isolates grew significantly better on glucose (Bayesian test) than on internal energy storage only, with norm ODmax values from 0.32 to 0.61. Ten of 11 isolates showed significant growth on HMW laminarin with norm ODmax values of 0.1 to 0.37, with the exception of FI145 (norm ODmax = 0). All isolates grew significantly better on partially hydrolyzed laminarin with norm ODmax values of 0.14 to 0.48 than on internal energy storage (Supplementary S2).

Species-specific significant (*p* < 0.05) difference in resource-specific growth among the 11 yeast isolates (p < 0.01; Kruskal-Wallis test) were observed (Supplementary S3).

### 3.3 Significant resource-specific growth of yeast isolates

The norm ODmax values of all isolates showed significant differences (*P* < 0.05; Kruskal-Wallis test) in resource specific growth on the three substrates, with the only exception of FI018 (Figure 2). The subsequent Dunńs test showed different growth on glucose (Bonferroni corrected *P* < 0.05; norm ODmax 0.36 - 0.61) than on HMW laminarin (norm ODmax 0 - 0.33) for nine of the isolates. Only for two isolates, FI121 (norm ODmax glucose: 0.36, HMW laminarin: 0.29) and FI018 (norm ODmax glucose: 0.32, HMW laminarin: 0.37), the pairwise comparison was not significant (Bonferroni corrected *P* > 0.05) (Figure 3, Supplementary S2).

**Figure 3:**
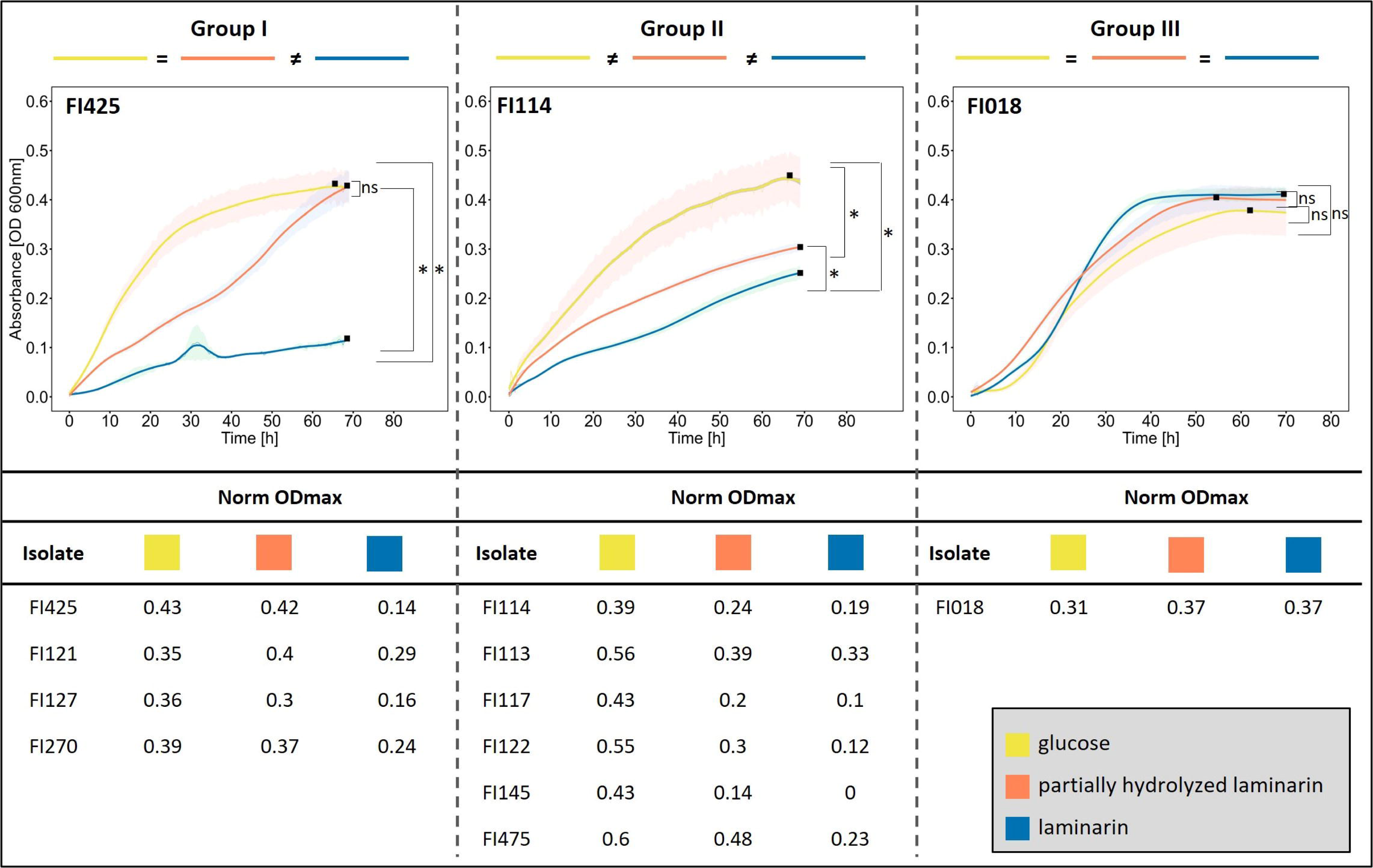
Yeasts were divided into three growth types. Group I, normalized OD maximum (filled square, Norm_ODmax) same for glucose and partially hydrolysed laminarin but significant difference to laminarin. Group II, Norm_ODmax significant difference across all substrates. Group III, Norm_ODmax same for all substrates. The figure shows the modelled growth curves for a representative isolate of each group. The shaded areas represent the standard error of the mean. Statistical comparisons between the three substrates were performed using the Kruskal-Wallis test and subsequent Dunn post hoc test with * for significant (p < 0.05) and ‘ns’ for non-significant differences.

Growth comparison on partially hydrolyzed laminarin (norm ODmax 0.14 – 0.48) versus glucose (norm ODmax 0.39 - 0.61) revealed a significance difference (Dunn’s test, Bonferroni corrected *P* < 0.05) for six isolates, namely FI113, FI114, FI117, FI122, FI145, FI475. The other five isolates (FI018, FI121, FI127, FI270 and FI425) grew the same on both substrates (Bonferroni corrected *P* > 0.05) (Figure 3, Supplementary S2).

Comparing growth on partially hydrolyzed laminarin versus HMW laminarin, all isolates performed significantly different (Dunn’s test, Bonferroni adjusted *P* < 0.05; norm ODmax partially hydrolyzed laminarin: 0.13 - 0.48, HMW laminarin: 0 - 0.33) except for FI018 (norm ODmax partially hydrolyzed laminarin: 0.36, HMW laminarin: 0.37) and FI113 (norm ODmax: 0.39 and 0.33, respectively) (Figure 3, Supplementary S3).

### 3.4 Tracking of laminarin degradation and intermediate product formation by FACE and PSA

While yeast isolates FI122, FI425 and FI475 moderately grew on the HMW precursor substrate laminarin (Figure 4 and Supplementary S4), they grew better on laminarin hydrolysate, as well as on laminarihexaose and laminaribiose. As revealed by PSA (quantitatively) and FACE (qualitatively), the HMW laminarin band gradually decreased in intensity across time points T0-T4, however, no intermediate degradation products were visible at any time point (Figure 3), suggesting that HMW laminarin was digested by yeast derived exo-acting glycosylhydrolysases. On the contrary, partially hydrolyzed laminarin was degraded via several intermediates ranging from laminari-pentaose, -tetraose, -triose and biose (Figure 3), characterized by a multitude of short-chained laminarin fragments at T0. As degradation progressed, the intensity of short-chained laminarin-based intermediates increased in favour of both, long-chained precursors and intermediates. To test if short laminarin-derived oligosaccharide precursors were indeed degraded via intermediate products each lacking one or several glucose units, the yeast isolates were incubated with pure laminarihexaose and -biose. FACE analysis revealed that the hexamer was gradually degraded by exo-laminarinase into intermediate degradation products of the pentaose and - tetraose- and -triose, biose and glucose (Figure 4). With ongoing temporal degradation, the intensity of the glucose band increased (Figure 4).

**Figure 4:**
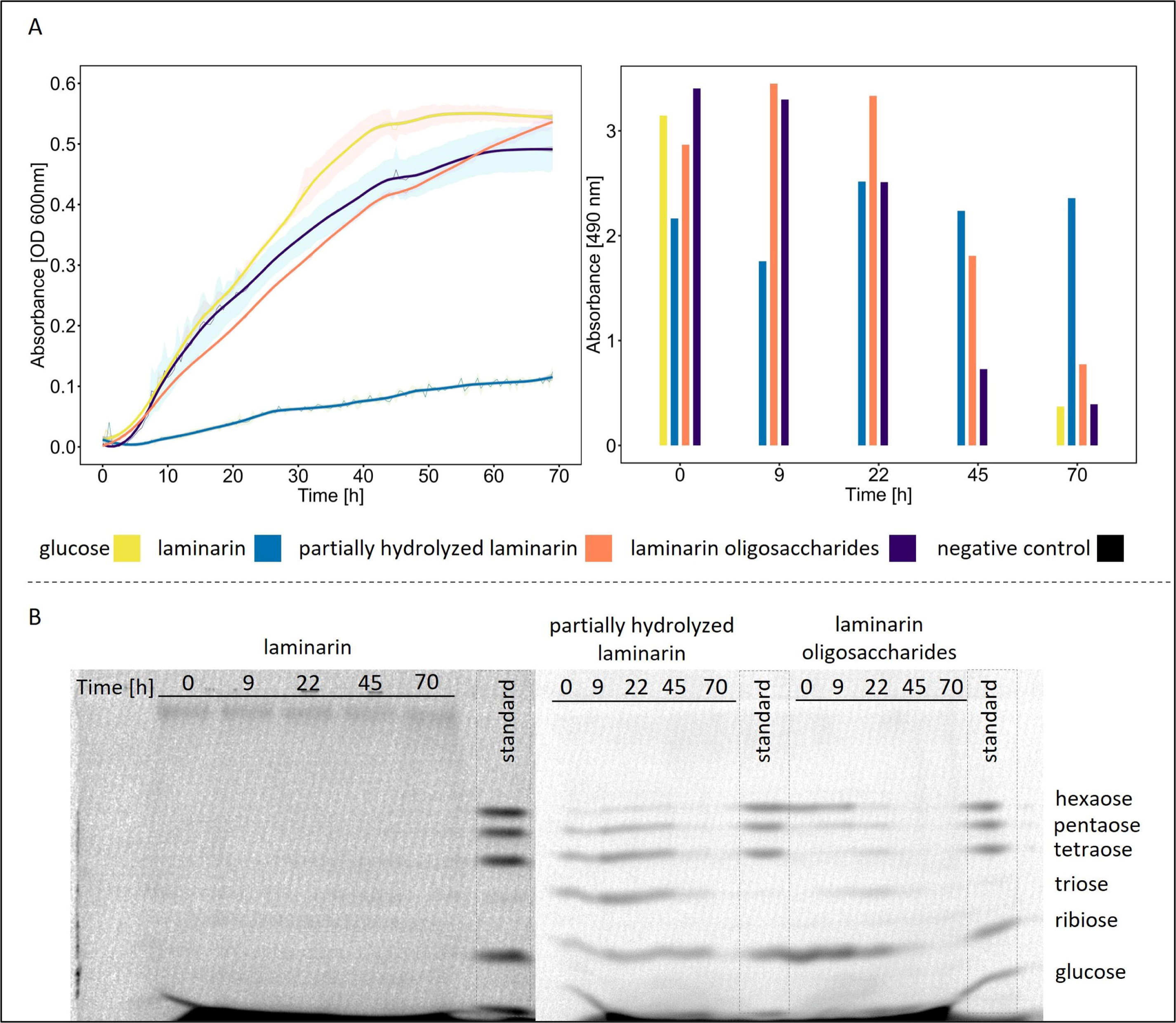
Tracking of laminarin-based degradation and intermediate products by fluorophore-assisted carbohydrate electrophoresis (FACE) and phenol-sulfuric acid analysis (PSA). A: The normalized growth (absorbance at 600 nm) of isolate FI475 on glucose, HMW laminarin (<10, >5 kDa), partially hydrolyzed laminarin (size unknown), and a mixture of two laminarin oligosaccharides over 70h with the shaded areas representing standard error of the mean (left), along with quantification by PSA (n = 1) (right). B: FACE precursor substrates and intermediate hydrolysis products at different time points. The dark black line at the bottom of the gel results from dye accumulation and partially coincides with the glucose band.

## 4 Discussion

Algal blooms composed of marine phytoplankton fix and convert atmospheric carbon dioxide into biological matter at the same magnitude as the combined global terrestrial plant biomass [41]. This organic matter is found in the key building blocks of eukaryotic algae and plants (organelles, energy storage, cell wall components). During the senescence stage of phytoplankton blooms, this material is released and bioavailable to microbial predators and scavengers at the base of the food web. Moreover, phytoplankton cells continuously release organic photosynthate molecules into the environment fueling their surrounding environment with organic matter, a large proportion of which are glycans. These large molecules are composed of linked monosaccharides and vary in a number of ways: the composition of different sugars, different functional groups on sugars, the linkage and branching pattern within defined repeating units, and the number of repeating units differentiating glycans into macromolecules of low and high molecular size and weight [42]. One of these glycans is laminarin, a central energy metabolite in microalgae. Almost 40% of organic carbon in particulate and dissolved organic matter is indeed made up of laminarin [24].

A globally important process in context of marine carbon cycling is the remineralization of phytoplankton-derived organic matter. It is now well described how algal blooms trigger secondary blooms of planktonic bacteria. Recurrent analyses of North Sea algal spring blooms revealed a structured succession of a specialized bacterial communities that sequentially degrade the complex mixture of algal polysaccharides during blooms [4, 43, 44]. Yet, recent studies lend increasing support to a role of marine fungi in organic carbon cycling. In the epipelagic zone, fungi reach ecologically relevant biomass levels [16, 45] which can temporally exceed the one of bacteria [15]. On particulate organic material in the bathypelagic zone, fungi are the most abundant microbial taxon [46]. Enzyme assays revealed alginase, carrageenase, fucoidanase and laminarinase activities in macroalgae-associated fungi [47]. SIP-analyses on natural pelagic microbial communities as well as fungal isolates, verified that marine fungi have the capacity to process phytoplankton-derived organic matter [19, 48]. Further, a global meta-transcriptomic study demonstrated the universal presence of fungal glycoside hydrolases, i.e., enzymes involved in glycan degradation, especially prevalent in ocean regions of high productivity [18]. In Priest et al. [16], we suggested that a markedly observed increase in yeast biomass towards the end of the spring phytoplankton bloom in Helgoland Roads was indicative of their high organic substrate degradation activity.

We developed a sensitive microtiter plate growth assay for marine yeasts, to (i) track substrate-induced growth in different yeast species in parallel at high temporal resolution, (ii) putative intermediate degradation products with the size of laminarin-based organic precursor substrates, and (iv) distinguish growth on intracellular storage products [49] from utilization of organic substrates, (see Supplementary 2 for details). Similar growth protocols enabling this analytical breadth are lacking (but see [20, 21, 50]. Given the prominence of laminarin as major phytoplankton bloom-derived glycan, this organic substrate as well as its fragments of lower molecular weight were deemed ecologically relevant proxies to study the degradation capacity of marine yeasts.

Observed growth patterns among eleven environmental yeast isolates differed with regard to species and organic substrate type. Generally, yeasts under investigation were divided into two categories that grew either the same or significantly less on HMW laminarin in comparison to partially hydrolyzed laminarin (Figure 1). Growth trajectories of yeasts belonging to the latter category were already distinguishable after 10 h post inoculation. A correlation to phylogeny or ecology of the yeast isolates was not observed. While partially hydrolyzed laminarin was composed of numerous shorter, but uncharacterized laminarin fragments (see Figure 3), similarly high growth kinetics were also achieved with a defined equimolar mixture of laminarihexaose and laminaribiose. While the HMW laminarin band gradually decreased in intensity over time without the appearance of intermediate degradation products (Figure 3), the degradation of partially hydrolyzed laminarin as well as of the laminarin oligosaccharide mix progressed via several intermediates ranging from laminari-pentaose, -tetraose, -triose and biose (Figure 3). With continuing degradation, the intensity of short-chained laminarin-based intermediates increased in favour of both, long-chained precursors and intermediates, demonstrating these yeasts utilized exo-acting glycosylhydrolysases. Together, these results demonstrate a wide spectrum of laminarin utilization among the yeasts under investigation, from direct and mostly slow degradation of HMW laminarin to faster growth and more complete utilization of laminarin-derived mono- and oligosaccharides.

Marine heterotrophic fungi have long been neglected as potential members and potential drivers of the microbial loop. Here we show for the first time that marine yeasts, given their observed substrate specificity, may occupy a specific niche in the microbial loop [51, 52], benefitting from bacterially derived carbohydrate degradation products [53]. This is supported by their documented appearance during the spring phytoplankton bloom at Helgoland Roads [16] (Figure 2). The proposed niche of marine yeasts in the microbial loop is congruent to the observed niche of terrestrial yeasts in upper forest soils [54]. Here, yeasts are efficient consumers of microbially derived (filamentous fungi and bacteria) carbohydrate degradation products [55]. Thus, yeasts appear to play a prominent role in microbial interactomes, both in marine and terrestrial ecosystems.

## 5 Acknowledgements

This project was funded as “05 Fokusprojekt 12-1” from the University of Bremen, Germany.

## 7 Data Availability Statement

All data generated in this project are provided in the form of tables and graphs in the main manuscript and Supplementary Data.

## 8 Authors contribution

MR, TH, JHH, AW, GG and MSJ conceived and designed the experiments. BAG, MSJ, TNH, and SN performed the experiments. BAG, TNH, RN, MSJ and SN analyzed the data. BAG, TH, and MR guided the data mining and wrote the paper. All authors have reviewed and agreed with the paper.

## 12 Supplementary Materials

Supplementary Material S1: Optimal growth parameters

Supplementary Material S2: Resource-specific growth of 11 yeast isolates on three substrates: glucose, HMW laminarin (<10, >5 kDa), and partially hydrolyzed laminarin (size unspecified). The figure displays growth curves and normalized maximum optical density (Norm_ODmax) for each isolate over a 70-hour period. Statistical comparisons between the three substrates were conducted using the Kruskal-Wallis test, followed by Dunn’s post hoc test. Results are denoted with an asterisk (*) for statistically significant differences (p < 0.05) and "ns" for non-significant differences.

Supplementary Material S3: Differences in resource-specific growth among 11 yeast isolates

Supplementary Material S4: Tracking of substrate degradation and intermediate products by fluorophore-assisted carbohydrate electrophoresis (FACE) and phenol-sulfuric acid (PSA) analysis

Supplementary Material S5: Quantification of glucose and laminarin-based precursor substrates

## References

1. Falkowski P. Ocean Science: The power of plankton. Nature. 2012;483:S17–20. doi:10.1038/483S17a.

2. Worden AZ, Follows MJ, Giovannoni SJ, Wilken S, Zimmerman AE, Keeling PJ. Environmental science. Rethinking the marine carbon cycle: factoring in the multifarious lifestyles of microbes. Science. 2015;347:1257594. doi:10.1126/science.1257594.

3. Azam, F., Fenchel, T., Field, J.G., Gray, J.S., Meyer-Reil, L.A., and Thingstad, F. The ecological role of water-column microbes in the sea. Marine Ecology Progress Series 10. 1983:257–63.

4. Teeling H, Fuchs BM, Bennke CM, Krüger K, Chafee M, Kappelmann L, et al. Recurring patterns in bacterioplankton dynamics during coastal spring algae blooms. Elife. 2016;5:e11888. doi:10.7554/eLife.11888.

5. Lee H, Bloxham B, Gore J. Resource competition can explain simplicity in microbial community assembly; 2023.

6. Lima-Mendez G, Faust K, Henry N, Decelle J, Colin S, Carcillo F, et al. Ocean plankton. Determinants of community structure in the global plankton interactome. Science. 2015;348:1262073. doi:10.1126/science.1262073.

7. Bjorbækmo MFM, Evenstad A, Røsæg LL, Krabberød AK, Logares R. The planktonic protist interactome: where do we stand after a century of research? ISME J. 2020;14:544–59. doi:10.1038/s41396-019-0542-5.

8. Sichert A, Corzett CH, Schechter MS, Unfried F, Markert S, Becher D, et al. Verrucomicrobia use hundreds of enzymes to digest the algal polysaccharide fucoidan. Nat Microbiol. 2020;5:1026–39. doi:10.1038/s41564-020-0720-2.

9. Reintjes G, Arnosti C, Fuchs B, Amann R. Selfish, sharing and scavenging bacteria in the Atlantic Ocean: a biogeographical study of bacterial substrate utilisation. ISME J. 2019;13:1119–32. doi:10.1038/s41396-018-0326-3.

10. Amend A, Burgaud G, Cunliffe M, Edgcomb VP, Ettinger CL, Gutiérrez MH, et al. Fungi in the Marine Environment: Open Questions and Unsolved Problems. mBio 2019. doi:10.1128/mBio.01189-18.

11. Peng X, Amend AS, Baltar F, Blanco-Bercial L, Breyer E, Burgaud G, et al. Planktonic Marine Fungi: A Review. JGR Biogeosciences 2024. doi:10.1029/2023JG007887.

12. Taylor JD, Cunliffe M. Multi-year assessment of coastal planktonic fungi reveals environmental drivers of diversity and abundance. ISME J. 2016;10:2118–28. doi:10.1038/ismej.2016.24.

13. Duan Y, Xie N, Song Z, Ward CS, Yung C-M, Hunt DE, et al. A High-Resolution Time Series Reveals Distinct Seasonal Patterns of Planktonic Fungi at a Temperate Coastal Ocean Site (Beaufort, North Carolina, USA). Appl Environ Microbiol 2018. doi:10.1128/AEM.00967-18.

14. Banos S, Gysi DM, Richter-Heitmann T, Glöckner FO, Boersma M, Wiltshire KH, et al. Seasonal Dynamics of Pelagic Mycoplanktonic Communities: Interplay of Taxon Abundance, Temporal Occurrence, and Biotic Interactions. Front Microbiol. 2020;11:1305. doi:10.3389/fmicb.2020.01305.

15. Gutiérrez MH, Pantoja S, Tejos E, Quiñones RA. The role of fungi in processing marine organic matter in the upwelling ecosystem off Chile. Mar Biol. 2011;158:205–19. doi:10.1007/s00227-010-1552-z.

16. Priest T, Fuchs B, Amann R, Reich M. Diversity and biomass dynamics of unicellular marine fungi during a spring phytoplankton bloom. Environ Microbiol. 2021;23:448–63. doi:10.1111/1462-2920.15331.

17. Drula E, Garron M-L, Dogan S, Lombard V, Henrissat B, Terrapon N. The carbohydrate-active enzyme database: functions and literature. Nucleic Acids Res. 2022;50:D571–D577. doi:10.1093/nar/gkab1045.

18. Chrismas N, Cunliffe M. Depth-dependent mycoplankton glycoside hydrolase gene activity in the open ocean-evidence from the Tara Oceans eukaryote metatranscriptomes. ISME J. 2020;14:2361–5. doi:10.1038/s41396-020-0687-2.

19. Orsi WD, Vuillemin A, Coskun ÖK, Rodriguez P, Oertel Y, Niggemann J, et al. Carbon assimilating fungi from surface ocean to subseafloor revealed by coupled phylogenetic and stable isotope analysis. ISME J. 2022;16:1245–61. doi:10.1038/s41396-021-01169-5.

20. Tamminen A, Happonen P, Barth D, Holmström S, Wiebe MG. High throughput, small scale methods to characterise the growth of marine fungi. PLoS One. 2020;15:e0236822. doi:10.1371/journal.pone.0236822.

21. Breyer E, Baltar F. The largely neglected ecological role of oceanic pelagic fungi. Trends Ecol Evol. 2023;38:870–88. doi:10.1016/j.tree.2023.05.002.

22. Diver P, Ward BA, Cunliffe M. Physiological and morphological plasticity in response to nitrogen availability of a yeast widely distributed in the open ocean. FEMS Microbiol Ecol 2024. doi:10.1093/femsec/fiae053.

23. Aluwihare LI, Repeta DJ. A comparison of the chemical characteristics of oceanic DOM and extracellular DOM produced by marine algae. Mar. Ecol. Prog. Ser. 1999;186:105–17. doi:10.3354/meps186105.

24. Becker S, Tebben J, Coffinet S, Wiltshire K, Iversen MH, Harder T, et al. Laminarin is a major molecule in the marine carbon cycle. Proc Natl Acad Sci U S A. 2020;117:6599– 607. doi:10.1073/pnas.1917001117.

25. Myklestad SM. Dissolved Organic Carbon from Phytoplankton. In: Abrajano TA, editor. The handbook of environmental chemistry. Berlin: Springer; 2000. p. 111–148. doi:10.1007/10683826_5.

26. Sidhu C, Kirstein IV, Meunier CL, Rick J, Fofonova V, Wiltshire KH, et al. Dissolved storage glycans shaped the community composition of abundant bacterioplankton clades during a North Sea spring phytoplankton bloom. Microbiome. 2023;11:77. doi:10.1186/s40168-023-01517-x.

27. Krause E, Wichels A, Erler R, Gerdts G. Study on the effects of near-future ocean acidification on marine yeasts: a microcosm approach. Helgol Mar Res. 2013;67:607–21. doi:10.1007/s10152-013-0348-1.

28. Berges JA, Franklin DJ, Harrison PJ. Evolution of an artifical seawater medium: Improvements in enriched seawater, artificial water over the last two decades. Journal of Phycology. 2001;37:1138–45. doi:10.1046/j.1529-8817.2001.01052.x.

29. Harrison PJ, Waters RE, Taylor FJR. A broad spectrum artificial sea water medium for coastal and open ocean phytoplankton. Journal of Phycology. 1980;16:28–35. doi:10.1111/j.0022-3646.1980.00028.x.

30. Wickerham J. Tropical Med. Hyg. 1939;42:176.

31. Reintjes G, Fuchs BM, Scharfe M, Wiltshire KH, Amann R, Arnosti C. Short-term changes in polysaccharide utilization mechanisms of marine bacterioplankton during a spring phytoplankton bloom. Environ Microbiol. 2020;22:1884–900. doi:10.1111/1462-2920.14971.

32. Redfield AC. The biological control of chemical factors in the environment. In: American Scientist 46(3). 1958;205–221.

33. Becker S, Scheffel A, Polz MF, Hehemann J-H. Accurate Quantification of Laminarin in Marine Organic Matter with Enzymes from Marine Microbes. Appl Environ Microbiol 2017. doi:10.1128/AEM.03389-16.

34. Ogura I, Sugiyama M, Tai R, Mano H, Matsuzawa T. Optimization of microplate-based phenol-sulfuric acid method and application to the multi-sample measurements of cellulose nanofibers; 2023.

35. Becker S, Hehemann J-H. Laminarin Quantification in Microalgae with Enzymes from Marine Microbes. Bio Protoc. 2018;8:e2666. doi:10.21769/BioProtoc.2666.

36. Jackson P. The use of polyacrylamide-gel electrophoresis for the high-resolution separation of reducing saccharides labelled with the fluorophore 8-aminonaphthalene- 1,3,6-trisulphonic acid. Detection of picomolar quantities by an imaging system based on a cooled charge-coupled device. Biochem J. 1990;270:705–13. doi:10.1042/bj2700705.

37. Midani FS, Collins J, Britton RA. AMiGA: Software for Automated Analysis of Microbial Growth Assays. mSystems. 2021;6:e0050821. doi:10.1128/mSystems.00508-21.

38. Tonner PD, Darnell CL, Engelhardt BE, Schmid AK. Detecting differential growth of microbial populations with Gaussian process regression. Genome Res. 2017;27:320–33. doi:10.1101/gr.210286.116.

39. Dinno A. dunn.test: Dunn’s Test of Multiple Comparisons Using Rank Sums: Version: 1.3.6. 2024-04-12. https://CRAN.R-project.org/package=dunn.test. Accessed 24 Jul 2024.

40. Wickham H, Chang W, Henry L, Pedersen TL, Takahashi K, Wilke C, et al. ggplot2: Elegant Graphics for Data Analysis. 2016. https://ggplot2.tidyverse.org. Accessed 12 Sep 2024.

41. Field CB, Behrenfeld MJ, Randerson JT, Falkowski P. Primary production of the biosphere: integrating terrestrial and oceanic components. Science. 1998;281:237–40. doi:10.1126/science.281.5374.237.

42. Arnosti C, Wietz M, Brinkhoff T, Hehemann J-H, Probandt D, Zeugner L; Amann, R. The Biogeochemistry of Marine Polysaccharides: Sources, Inventories, and Bacterial Drivers of the Carbohydrate Cycle. Ann Rev Mar Sci. 2021;13:81–108. doi:10.1146/annurev-marine-032020-012810.

43. Chafee M, Fernàndez-Guerra A, Buttigieg PL, Gerdts G, Eren AM, Teeling H; Amann, Rudolf I. Recurrent patterns of microdiversity in a temperate coastal marine environment. ISME J. 2018;12:237–52. doi:10.1038/ismej.2017.165.

44. Lucas J, Wichels A, Teeling H, Chafee M, Scharfe M, Gerdts G. Annual dynamics of North Sea bacterioplankton: seasonal variability superimposes short-term variation. FEMS Microbiol Ecol. 2015;91:fiv099. doi:10.1093/femsec/fiv099.

45. Hassett BT, Borrego EJ, Vonnahme TR, Rämä T, Kolomiets MV, Gradinger R. Arctic marine fungi: biomass, functional genes, and putative ecological roles. ISME J. 2019;13:1484–96. doi:10.1038/s41396-019-0368-1.

46. Bochdansky AB, Clouse MA, Herndl GJ. Eukaryotic microbes, principally fungi and labyrinthulomycetes, dominate biomass on bathypelagic marine snow. ISME J. 2017;11:362–73. doi:10.1038/ismej.2016.113.

47. Patyshakuliyeva A, Falkoski DL, Wiebenga A, Timmermans K, Vries RP de. Macroalgae Derived Fungi Have High Abilities to Degrade Algal Polymers. Microorganisms 2019. doi:10.3390/microorganisms8010052.

48. Cunliffe M, Hollingsworth A, Bain C, Sharma V, Taylor JD. Algal polysaccharide utilisation by saprotrophic planktonic marine fungi. Fungal Ecology. 2017;30:135–8. doi:10.1016/j.funeco.2017.08.009.

49. Athenaki M, Gardeli C, Diamantopoulou P, Tchakouteu SS, Sarris D, Philippoussis A; Papanikolaou, S. Lipids from yeasts and fungi: physiology, production and analytical considerations. J Appl Microbiol. 2018;124:336–67. doi:10.1111/jam.13633.

50. Pang K-L, Chiang MW-L, Guo S-Y, Shih C-Y, Dahms HU, Hwang J-S; Cha, Hyo-Jung. Growth study under combined effects of temperature, pH and salinity and transcriptome analysis revealed adaptations of Aspergillus terreus NTOU4989 to the extreme conditions at Kueishan Island Hydrothermal Vent Field, Taiwan. PLoS One. 2020;15:e0233621. doi:10.1371/journal.pone.0233621.

51. Francis B, Urich T, Mikolasch A, Teeling H, Amann R. North Sea spring bloom-associated Gammaproteobacteria fill diverse heterotrophic niches. Environ Microbiome. 2021;16:15. doi:10.1186/s40793-021-00385-y.

52. Krüger K, Chafee M, Ben Francis T, Del Glavina Rio T, Becher D, Schweder T, et al. In marine Bacteroidetes the bulk of glycan degradation during algae blooms is mediated by few clades using a restricted set of genes. ISME J. 2019;13:2800–16. doi:10.1038/s41396-019-0476-y.

53. Krabberød AK, Bjorbækmo MF, Shalchian-Tabrizi K, Logares R. Exploring the oceanic microeukaryotic interactome with metaomics approaches. Aquat. Microb. Ecol. 2017;79:1–12. doi:10.3354/ame01811.

54. Martinović T, Mašínová T, López-Mondéjar R, Jansa J, Štursová M, Starke R; Baldrian, Petr. Microbial utilization of simple and complex carbon compounds in a temperate forest soil. Soil Biology and Biochemistry. 2022;173:108786. doi:10.1016/j.soilbio.2022.108786.

55. Mašínová T, Yurkov A, Baldrian P. Forest soil yeasts: Decomposition potential and the utilization of carbon sources. Fungal Ecology. 2018;34:10–9. doi:10.1016/j.funeco.2018.03.005.

56. Niggemeier S. Created in BioRender.: Subscription: Institution - Academic Agreement number: CV27APCPD6. 2024. BioRender.com/o73b845.

