## Supplemental Material for "Utilization and degradation of laminarin-based substrates by marine yeasts suggests their niche-specific role in microbial loop dynamics"

### Supplementary Information

*Corresponding author

Marlis Reich, University of Bremen, FB2, Molecular Ecology,

Leobener Str. 6, 28359 Bremen, Germany

**Supplementary material 1: Optimal growth parameters**

Significant values in Bayesian test are shown with an asterisk (*), indicating that the growth on this substrate was significantly different from growth of internal storage products. Only significant comparisons (Bayesian test) were analyzed by Kruskal-Wallis (K-W) followed by Dunn's test. The test results are represented with *p* values and "ns" indicates non-significant results.

**Supplementary material 1.A: $NaNO_3$ (N) and $NaH_2PO_4$ (P) concentrations**

(A) All isolates showed significantly (Dunn's test, Bonferroni corrected P < 0.01) higher norm ODmax in medium containing 75 µM and 30 µM $NaNO_3$ (N) and $NaH_2PO_4$ (P) (norm ODmax 0.26 – 0.33) compared to 750 µM and 300 µM $NaNO_3$ and $NaH_2PO_4$ (norm ODmax 0 – 0.12), respectively.

| Isolates | Substrate | Norm ODmax |
|---|---|---|
| FI018 | 750umN_300umP | 0.12165* |
| FI114 | 750umN_300umP | 0.03725* |
| FI117 | 750umN_300umP | -0.01165* |
| FI018 | 75umN_30umP | 0.331675* |
| FI114 | 75umN_30umP | 0.269625* |
| FI117 | 75umN_30umP | 0.264625* |

| Isolates | Substrate comparison | K-W test | Dunn's test |
|---|---|---|---|
| FI018 | 750umN_300umP vs. 75umN_30umP | *p = 0.02092* | *p = 0.01046* |
| FI114 | 750umN_300umP vs. 75umN_30umP | *p = 0.02092* | *p = 0.01046* |
| FI117 | 750umN_300umP vs. 75umN_30umP | *p = 0.02092* | *p = 0.01046* |

**Supplementary material 1.B: Redfield stoichiometry**

(B) All isolates grew the same (Dunn's test, Bonferroni corrected P > 0.05) at Redfield stoichiometry of C:N:P of 106:16:1 (norm ODmax 0.48 - 0.61) and C:N:P of 173:16:1 (norm ODmax 0.46 - 0.6) with high carbon (glucose) concentrations. However, at a Redfield stoichiometry of C:N:P of 106:16:1, the observed growth significantly differed from the C:N:P ratio of 17.3:16:1 (norm ODmax 0.19 – 0.29) at low carbon (glucose) concentrations (Dunn's test, Bonferroni corrected P < 0.02).

Two controls with N and P concentrations as in (A) and stoichiometric ratios of C:N:P of 17.3:2.5:1 and 173:2.5:1, respectively, identified carbon as the main driver of growth. For the high carbon (glucose) control at a C:N:P ratio of 173:2.5:1 (norm ODmax 0.43 – 0.63), no isolate showed a significant difference in growth compared to the C:N:P ratio of 173:16:1 (norm Odmax 0.46 – 0.6). For the low carbon control at C:N:P of 17.3:2.5:1, only isolate FI113 showed significant different growth (Dunn's test, Bonferroni corrected $P < 0.05$) with norm ODmax of 0.17 compared to C:N:P of 17.3:16:1 with a norm ODmax of 0.19.

| Isolates | Substrate | C:N:P ratio | Norm_ODmax |
|---|---|---|---|
| FI113 | Redfield | 106:16:01 | 0.6133* |
| FI114 | Redfield | 106:16:01 | 0.480225* |
| FI117 | Redfield | 106:16:01 | 0.55235* |
| FI113 | Redfield_high_glu | 173.3:16:1 | 0.6077* |
| FI114 | Redfield_high_glu | 173.3:16:1 | 0.459275* |
| FI117 | Redfield_high_glu | 173.3:16:1 | 0.4756* |
| FI113 | Redfield_low_glu | 17.333:16:1 | 0.20585* |
| FI114 | Redfield_low_glu | 17.333:16:1 | 0.256175* |
| FI117 | Redfield_low_glu | 17.333:16:1 | 0.1948* |
| FI113 | Control_high | 173.3:2.5:1 | 0.62605* |
| FI114 | Control_high | 173.3:2.5:1 | 0.426475* |
| FI117 | Control_high | 173.3:2.5:1 | 0.473775* |
| FI113 | Control_low | 17.33:2.5:1 | 0.23885* |
| FI114 | Control_low | 17.33:2.5:1 | 0.1822* |
| FI117 | Control_low | 17.33:2.5:1 | 0.168925* |

| Isolates | Substrate comparison | K-W test | Dunn's test |
|---|---|---|---|
| FI113 | Redfield vs. Redfield_high | *p = 0.0066* | ns |
| FI114 | Redfield vs. Redfield_high | *p = 0.00582* | ns |
| FI117 | Redfield vs. Redfield_high | *p = 0.00747* | ns |
| FI113 | Redfield vs. Redfield_low | *p = 0.0066* | *p = 0.0104* |

| | | | |
|---|---|---|---|
| FI114 | Redfield vs. Redfield_low | *p = 0.00582* | *p = 0.0104* |
| FI117 | Redfield vs. Redfield_low | *p = 0.00747* | *p = 0.0104* |
| FI113 | Redfield_high vs. Control_high | *p = 0.0066* | ns |
| FI114 | Redfield_high vs. Control_high | *p = 0.00582* | ns |
| FI117 | Redfield_high vs. Control_high | *p = 0.00747* | ns |
| FI113 | Redfield_low vs. Control_low | *p = 0.0066* | *p = 0.04163* |
| FI114 | Redfield_low vs. Control_low | *p = 0.00582* | ns |
| FI117 | Redfield_low vs. Control_low | *p = 0.00747* | ns |

**Supplementary material 1.C: Growth in ASW vs. EASW medium**

(C) All isolates grew to a significantly higher norm ODmax (Dunn's test, Bonferroni corrected P < 0.02; glucose: 0.39 – 0.56; HMW laminarin: 0.1 – 0.33) at pH 7.0 (ASW) than at pH 8.2 (EASW, norm ODmax glucose: 0.17 – 0.39; laminarin: 0.02 – 0.04), regardless of the carbon source.

| Isolates | Substrate | Norm ODMAX |
|---|---|---|
| FI113 | Glucose_ASW | 0.563875* |
| FI114 | Glucose_ASW | 0.393925* |
| FI117 | Glucose_ASW | 0.435025* |
| FI113 | Glucose_EASW | 0.3938* |
| FI114 | Glucose_EASW | 0.171975* |
| FI117 | Glucose_EASW | 0.279225* |
| FI113 | Laminarin_ASW | 0.33125* |
| FI114 | Laminarin_ASW | 0.18895* |
| FI117 | Laminarin_ASW | 0.10245* |
| FI113 | Laminarin_EASW | 0.0406* |
| FI114 | Laminarin_EASW | 0.0181* |
| FI117 | Laminarin_EASW | 0.0288* |

| Isolates | Substrate comparison | K-W test | Dunn's test |
|---|---|---|---|
| FI113 | Glucose_ASW vs. Glucose_EASW | *p = 0.02749* | *p = 0.01046* |
| FI114 | Glucose_ASW vs. Glucose_EASW | *p = 0.04111* | *p = 0.01046* |
| FI117 | Glucose_ASW vs. Glucose_EASW | *p = 0.02749* | *p = 0.01046* |
| FI113 | Laminarin_ASW vs. Laminarin_EASW | *p = 0.02749* | *p = 0.01046* |
| FI114 | Laminarin_ASW vs. Laminarin_EASW | *p = 0.04111* | *p = 0.01046* |
| FI117 | Laminarin_ASW vs. Laminarin_EASW | *p = 0.02749* | *p = 0.01046* |

**Supplementary material 1.D: Nutrient deprivation times**

(D) Nutrient deprivation prior to culture experiments had a significant effect on growth (Dunn's test, Bonferroni corrected P < 0.01). However, the effect differed among isolates. FI114 and FI117 grew significantly better without nutrient deprivation (norm ODmax 0.47 - 0.53) for 45h (norm ODmax 0.27 - 0.29) and 65h (norm ODmax 0.33 - 0.34). In contrast, FI113 showed higher growth with deprivation of 45h (norm ODmax 0.43) and 65h (norm ODmax 0.63) compared to those without nutrient deprivation (norm ODmax 0.26).

| Isolates | Substrate | Norm_ODMAX |
|---|---|---|
| FI113 | No-deprivation | 0.264225* |
| FI114 | No-deprivation | 0.4698* |
| FI117 | No-deprivation | 0.524225* |
| FI113 | 45h-deprivation | 0.428725* |
| FI114 | 45h-deprivation | 0.269625* |
| FI117 | 45h-deprivation | 0.289625* |
| FI113 | 65h-deprivation | 0.62605* |
| FI114 | 65h-deprivation | 0.326175* |
| FI117 | 65h-deprivation | 0.338825* |

| Isolates | Substrate comparison | K-W test | Dunn's test |
|---|---|---|---|
| FI113 | No-deprivation vs. 45h-deprivation | *p = 0.0097* | *p = 0.021654* |
| FI114 | No-deprivation vs. 45h-deprivation | *p = 0.0154* | *p = 0.00668* |
| FI117 | No-deprivation vs. 45h-deprivation | *p = 0.01246* | *p = 0.004896* |
| FI113 | No-deprivation vs. 65h-deprivation | *p = 0.0097* | *p = 0.01046* |
| FI114 | No-deprivation vs. 65h-deprivation | *p = 0.0154* | *p = 0.01046* |
| FI117 | No-deprivation vs. 65h-deprivation | *p = 0.01246* | *p = 0.01046* |

**Supplementary material 1.E: Growth detection limits on glucose and laminarin**

(E) All isolates grew significantly different on glucose compared to the negative control (Bayesian test). Growth was negatively correlated with glucose concentration, albeit at 25 g/l glucose growth (norm ODmax 0.21 - 0.56) was negatively affected and lower than at 2.5 g/L (norm ODmax 0.32 – 0.77). The detection limit for fungal growth on glucose differed among the isolates. Isolates FI114 and FI117 did not grow below 0.25 g/l glucose (norm ODmax of 0.13 and 0.14). The growth of isolate FI018 was detectable, even at 0.00025 g/l glucose (ODmax 0.07).

The lowest HMW laminarin concentration employed (0.017 g/l) did not result in significant differences in the normalized ODmax of any of the isolates compared to growth on internal energy storage (Bayesian test). Consequently, this concentration was not considered in the subsequent experiments. Growth on 0.17 g/l HMW laminarin was significant only for isolate FI117 (ODmax 0.18). Across all three isolates, significant growth was observed only at the highest HMW laminarin concentration of 0.97 g/L (norm ODmax 0.8-0.26).

| Isolates | Substrate | Norm_ODMAX |
|---|---|---|
| FI018 | Glucose_25g/L | 0.56425* |
| FI114 | Glucose_25g/L | 0.2137* |
| FI117 | Glucose_25g/L | 0.21955* |
| FI018 | Glucose_2.5g/L | 0.7759* |
| FI114 | Glucose_2.5g/L | 0.326175* |
| FI117 | Glucose_2.5g/L | 0.338825* |
| FI018 | Glucose_0.25g/L | 0.2464* |
| FI114 | Glucose_0.25g/L | 0.130425* |

| | | |
|---|---|---|
| FI117 | Glucose_0.25g/L | 0.11775* |
| FI018 | Glucose_0.025g/L | 0.051725* |
| FI114 | Glucose_0.025g/L | -0.00137 |
| FI117 | Glucose_0.025g/L | -0.02598 |
| FI018 | Glucose_0.0025g/L | 0.044425* |
| FI114 | Glucose_0.0025g/L | 0.0124 |
| FI117 | Glucose_0.0025g/L | -0.03173 |
| FI018 | Glucose_0.00025g/L | 0.225575* |
| FI114 | Glucose_0.00025g/L | 0.015575 |
| FI117 | Glucose_0.00025g/L | -0.041 |
| FI018 | Laminarin_0.17g/L | 0.00405 |
| FI114 | Laminarin_0.17g/L | 0.01325 |
| FI117 | Laminarin_0.17g/L | 0.025275* |
| FI018 | Laminarin_0.97g/L | 0.2583* |
| FI114 | Laminarin_0.97g/L | 0.178* |
| FI117 | Laminarin_0.97g/L | 0.08375* |

**Supplementary material 1.F: Effect of priming during nutrient deprivation**

(F) Isolates FI113, FI114 and FI117 grew better on HMW laminarin without priming during the pre-culture phase. This was significantly different only for isolate FI113 (Dunn's test, Bonferroni corrected $P < 0.01$). The norm ODmax values of the unprimed isolates were 0.15–0.21, while those of the primed isolates were 0.04–0.18.

| Isolates | Substrate | Norm_ODMAX |
|---|---|---|
| FI113 | Unprimed_laminarin | 0.151806* |
| FI114 | Unprimed_laminarin | 0.176981* |
| FI117 | Unprimed_laminarin | 0.209881* |
| FI113 | Primed_laminarin | 0.038725* |
| FI114 | Primed_laminarin | 0.14295* |
| FI117 | Primed_laminarin | 0.182775* |

| Isolates | Substrate comparison | K-W test | Dunn's test |
|---|---|---|---|
| FI113 | Unprimed vs. Primed_laminarin | *p = 0.02092* | *p = 0.01046* |
| FI114 | Unprimed vs. Primed_laminarin | ns | ns |
| FI117 | Unprimed vs. Primed_laminarin | ns | ns |

**Supplementary material 2: Resource-specific growth of 11 yeast isolates on three substrates: glucose, HMW laminarin (<10, >5 kDa), and partially hydrolyzed laminarin (size unspecified)**

The figure displays growth curves and normalized maximum optical density (Norm_ODmax) for each isolate over a 70-hour period. Statistical comparisons between the three substrates were conducted using the Kruskal-Wallis test, followed by Dunn's post hoc test. Results are denoted with an asterisk (*) for statistically significant differences (p < 0.05) and "ns" for non-significant differences.

#### *Supplementary material 2*

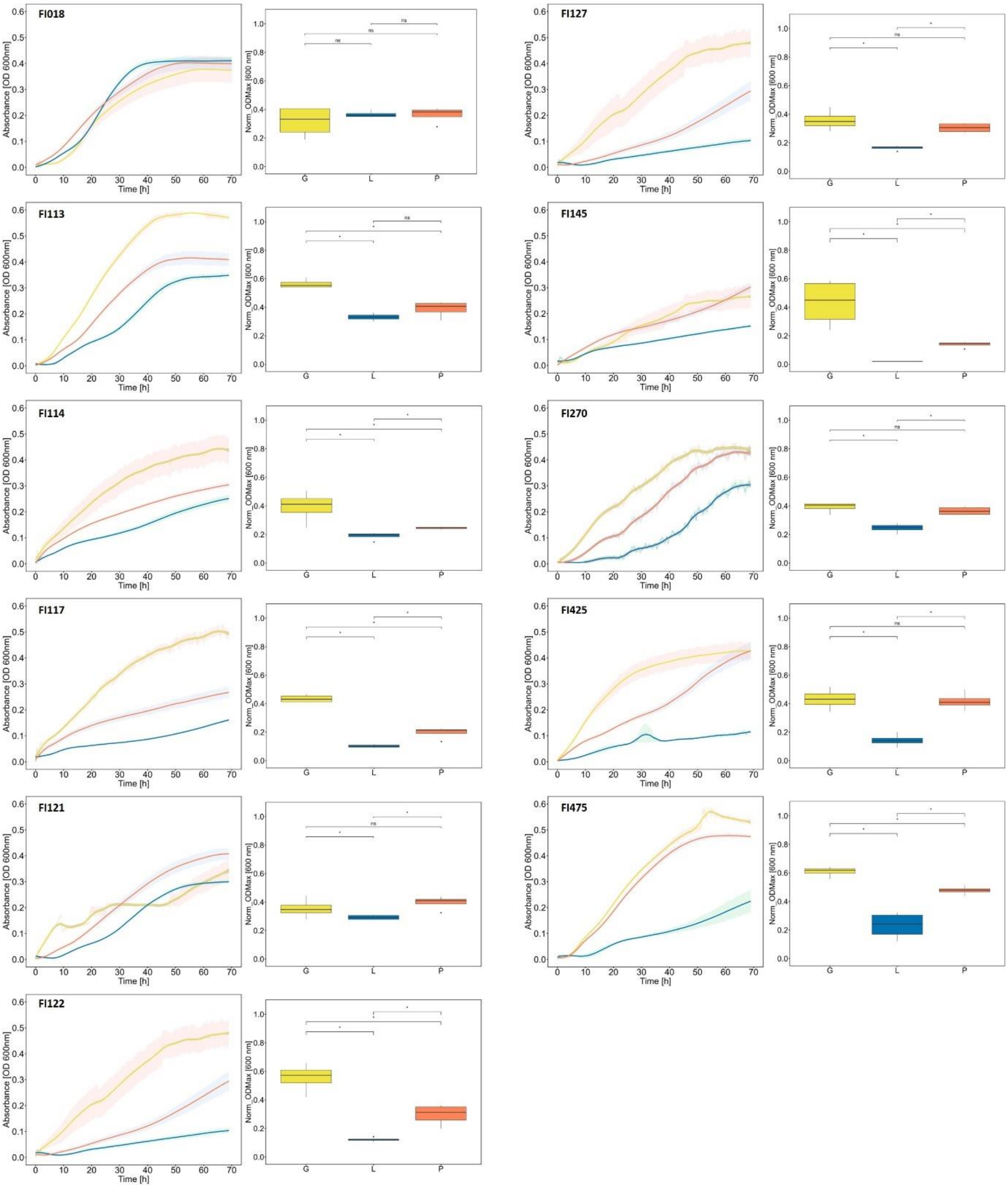

| | Glucose vs. partially hydrolyzed laminarin | Glucose vs. laminarin | Partially hydrolyzed laminarin vs. laminarin |
|---|---|---|---|
| FI018 | ns | ns | ns |
| FI113 | *0.011* | *0.011* | ns |
| FI114 | *0.011* | *0.041* | *0.011* |
| FI117 | *0.011* | *0.011* | *0.011* |
| FI121 | *0.041* | ns | *0.011* |
| FI122 | *0.011* | *0.011* | *0.011* |
| FI127 | *0.011* | ns | *0.011* |
| FI145 | *0.011* | *0.011* | *0.011* |
| FI270 | *0.011* | ns | *0.011* |
| FI425 | *0.011* | ns | *0.011* |
| FI475 | *0.011* | *0.011* | *0.011* |

**Supplementary material 3: Differences in resource-specific growth among 11 yeast isolates**

Significant values (Bayesian test) carry an asterisk (*), indicating that growth on this substrate was significant in comparison to growth of internal storage.

| Isolates | Substrate | Norm_ODMAX |
|---|---|---|
| FI018 | Glucose | 0.31525* |
| FI113 | Glucose | 0.563875* |
| FI114 | Glucose | 0.393925* |
| FI117 | Glucose | 0.435025* |
| FI121 | Glucose | 0.355183* |
| FI122 | Glucose | 0.555166* |

| | | |
|---|---|---|
| FI127 | Glucose | 0.356525* |
| FI145 | Glucose | 0.431775* |
| FI270 | Glucose | 0.390575* |
| FI425 | Glucose | 0.431375* |
| FI475 | Glucose | 0.608725* |
| FI018 | Laminarin | 0.365975* |
| FI113 | Laminarin | 0.33125* |
| FI114 | Laminarin | 0.18895* |
| FI117 | Laminarin | 0.10245* |
| FI121 | Laminarin | 0.29235* |
| FI122 | Laminarin | 0.12245* |
| FI127 | Laminarin | 0.163475* |
| FI145 | Laminarin | -0.01525* |
| FI270 | Laminarin | 0.2449* |
| FI425 | Laminarin | 0.14225* |
| FI475 | Laminarin | 0.23215* |
| FI018 | Par. dig. laminarin | 0.36625* |
| FI113 | Par. dig. laminarin | 0.3883* |
| FI114 | Par. dig. laminarin | 0.245425* |
| FI117 | Par. dig. laminarin | 0.196825* |
| FI121 | Par. dig. laminarin | 0.395925* |
| FI122 | Par. dig. laminarin | 0.296375* |
| FI127 | Par. dig. laminarin | 0.304625* |
| FI145 | Par. dig. laminarin | 0.13855* |
| FI270 | Par. dig. laminarin | 0.365675* |
| FI425 | Par. dig. laminarin | 0.416875* |
| FI475 | Par. dig. laminarin | 0.477175* |

The growth of isolates was evaluated separately with glucose, HMW laminarin, and partially hydrolyzed laminarin. Only significant values (Bayesian test) were analyzed further. The Kruskal-Wallis test revealed significant differences across all substrates, with p-values of 0.0033, 0.0003, and 0.0002. Dunn’s post hoc test was applied, assigning p-values to each substrate. “ns” indicates non-significant results.

For glucose, the subsequent Dunn's test revealed for one pairwise comparison a significant (Bonferroni adjusted P < 0.04) diverging growth performance, namely between the most performing (FI475, norm ODmax = 0.61) and the worst performing isolate (FI018, norm ODmax = 0.32). The significant differences were observed between most performing isolates (FI113, FI122, FI475) and worse performing isolates (FI018, FI121, FI127) on glucose.

<u>Glucose</u>

| | FI018 | FI113 | FI114 | FI117 | FI121 | FI122 | FI127 | FI145 | FI270 | FI425 | FI475 |
|---|---|---|---|---|---|---|---|---|---|---|---|
| FI018 | - | | | | | | | | | | |
| FI113 | *0.011* | - | | | | | | | | | |
| FI114 | ns | *0.011* | - | | | | | | | | |
| FI117 | *0.011* | *0.011* | ns | - | | | | | | | |
| FI121 | ns | *0.011* | ns | *0.042* | - | | | | | | |
| FI122 | *0.011* | ns | *0.042* | *0.042* | *0.022* | - | | | | | |
| FI127 | ns | *0.011* | ns | *0.042* | ns | *0.022* | - | | | | |
| FI145 | ns | ns | ns | ns | ns | ns | ns | - | | | |
| FI270 | ns | *0.011* | ns | ns | ns | *0.011* | ns | ns | - | | |
| FI425 | *0.011* | ns | *0.042* | ns | *0.022* | ns | *0.022* | ns | *0.042* | - | |
| FI475 | *0.011* | *0.042* | *0.011* | ns | *0.011* | *0.011* | *0.011* | *0.011* | *0.011* | ns | - |

For resource-specific growth on HMW laminarin, highest amount of pairwise comparisons with the two isolates FI018 and FI113 were identified as significantly different (Dunn's test, Bonferroni adjusted $P < 0.04$). FI018 was the isolate with the highest norm ODmax value (0.37) on laminarin and showed significant differences compared to the norm ODmax of all isolates except of FI425. Similarly, FI113, which exhibited the second-highest norm ODmax of 0.33 on HMW laminarin, demonstrated a significant difference from all other isolates except of FI425. While with the lowest values of norm ODmax, which were 0.1, 0.12, found in FI117, FI122, respectively.

<u>HMW Laminarin</u>

| | FI018 | FI113 | FI114 | FI117 | FI121 | FI122 | FI127 | FI145 | FI270 | FI425 | FI475 |
|---|---|---|---|---|---|---|---|---|---|---|---|
| FI018 | - | | | | | | | | | | |
| FI113 | *0.042* | - | | | | | | | | | |
| FI114 | *0.011* | *0.011* | - | | | | | | | | |
| FI117 | *0.011* | *0.011* | *0.011* | - | | | | | | | |
| FI121 | *0.011* | *0.042* | *0.011* | *0.011* | - | | | | | | |
| FI122 | *0.011* | *0.011* | *0.011* | *0.022* | *0.011* | - | | | | | |
| FI127 | *0.011* | *0.011* | ns | *0.011* | *0.011* | *0.022* | - | | | | |
| FI145 | *0.011* | *0.011* | *0.011* | *0.011* | *0.011* | *0.011* | *0.011* | - | | | |
| FI270 | *0.011* | *0.011* | *0.042* | *0.011* | *0.022* | *0.011* | *0.011* | *0.011* | - | | |
| FI425 | ns | ns | ns | ns | ns | ns | ns | ns | ns | - | |
| FI475 | *0.011* | *0.042* | ns | *0.011* | ns | *0.042* | ns | *0.011* | ns | ns | - |

A significant difference (Dunn's test, Bonferroni adjusted P < 0.05) in growth on partially hydrolyzed laminarin was observed among many isolates. The two isolates FI475 and FI425, which exhibited highest normalized ODmax values (0.48; 0.42), with the two isolates FI145 and FI117 demonstrating the smallest normalized ODmax values (0.14; 0.2).

Partially hydrolyzed laminarin

| | FI018 | FI113 | FI114 | FI117 | FI121 | FI122 | FI127 | FI145 | FI270 | FI425 | FI475 |
|---|---|---|---|---|---|---|---|---|---|---|---|
| FI018 | - | | | | | | | | | | |
| FI113 | *0.011* | - | | | | | | | | | |
| FI114 | *0.011* | *0.011* | - | | | | | | | | |
| FI117 | *0.011* | *0.011* | *0.011* | - | | | | | | | |
| FI121 | ns | ns | *0.011* | *0.011* | - | | | | | | |
| FI122 | *0.042* | *0.042* | ns | ns | *0.042* | - | | | | | |
| FI127 | ns | *0.042* | *0.011* | *0.011* | *0.042* | ns | - | | | | |
| FI145 | *0.011* | ns | *0.011* | ns | *0.011* | *0.011* | *0.011* | - | | | |
| FI270 | ns | ns | *0.011* | *0.011* | ns | ns | *0.011* | *0.011* | - | | |
| FI425 | ns | ns | *0.011* | *0.011* | ns | *0.042* | *0.042* | *0.011* | ns | - | |
| FI475 | *0.011* | *0.011* | *0.011* | *0.011* | *0.022* | *0.011* | *0.011* | *0.011* | *0.011* | ns | - |

**Supplementary material 4: Tracking of substrate degradation and intermediate products by fluorophore-assisted carbohydrate electrophoresis (FACE) and phenol-sulfuric acid (PSA) analysis**

To track the degradation of laminarin-based precursor substrates over time and identify intermediate degradation products, samples were taken at five time points (0, 9, 22, 45, and 70 h) and profiled with FACE or quantified by PSA (n =1) using three yeast isolates, FI475 (See Results Fig.3), FI122, and FI425. In addition to the HMW laminarin and partially hydrolyzed laminarin, a mixture of two laminarin oligosaccharides (laminarihexaose and laminaribiose) was used in the standard growth protocol.

The top left section displays the normalized growth (absorbance at 600 nm) of isolates (grown in replication of n = 4) FI122 and FI425 on glucose, HMW laminarin (<10, >5 kDa), partially hydrolyzed laminarin (size unknown), and a mixture of two laminarin oligosaccharides over 70h, along with quantification by PSA (top right). The shaded areas represent standard error of the mean. The FACE gel in the lower section shows the precursor substrate and its intermediate hydrolysis products at various time points. The dark black line at the bottom of the gel results from dye accumulation and partially coincides with the glucose band.

The isolates showed enhanced growth on partially hydrolyzed laminarin and laminarin oligosaccharides compared to the high-molecular-weight (HMW) laminarin precursor. This was reflected in the increased intensity of bands corresponding to intermediate products from the low-molecular-weight laminarin and oligosaccharides over time (0-70 h). Conversely, no intermediate degradation products were detected for HMW laminarin at any time point, suggesting that it was digested by exo-acting glycosylhydrolases produced by yeast.

#### *Supplementary material 4*

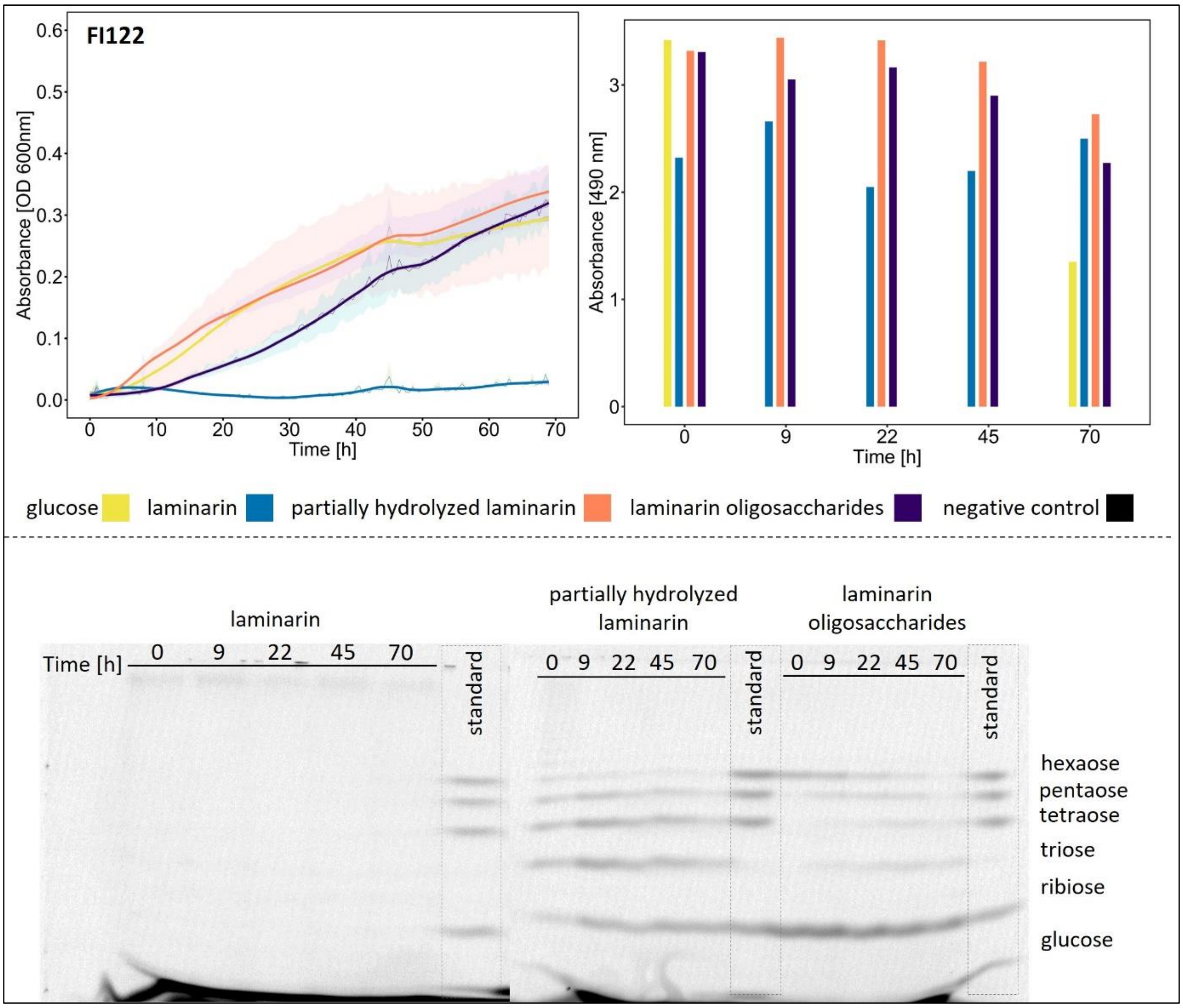

##### *Supplementary material 4, continued*

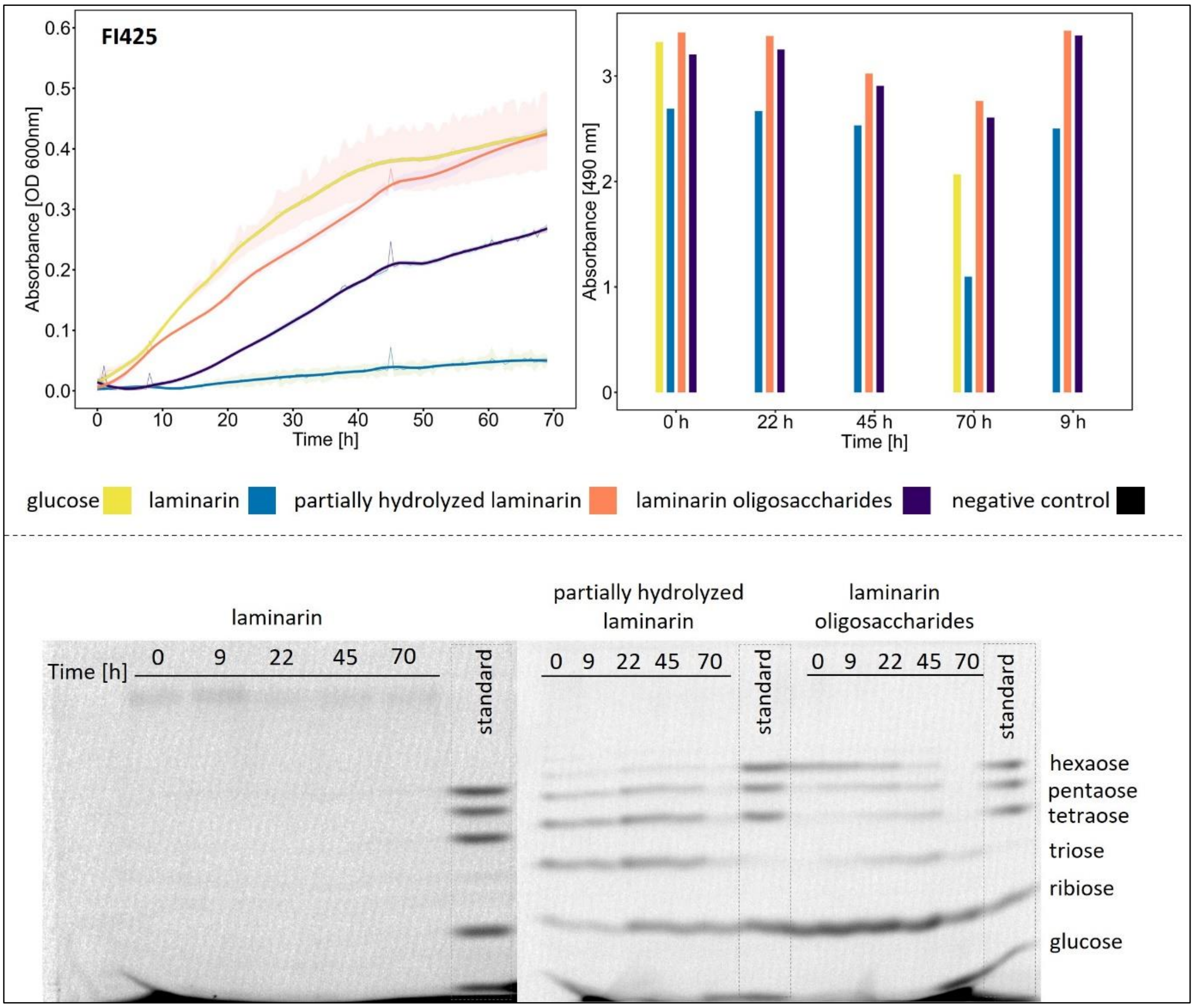

**Supplementary material 5: Quantification of glucose and laminarin-based precursor substrates**

The growth of 11 yeast isolates on HMW laminarin (<10, >5 kDa) and partially hydrolyzed laminarin (size unknown) as the sole carbon source was compared, using a normalized concentration of 0.85 g/l glucose equivalents (PSA, see Material and Methods 2.4) corresponding to 0.97 g/l HMW laminarin (estimated repeating unit of n = 10), The positive control contained 0.85 g/l glucose, the negative control contained ASW medium only. Samples were collected at five time points (0, 9, 22, 45, and 70 h) and analyzed by PSA (n = 1). The negative controls for FI121 at 70 hours and FI122 at 0 hours could not be quantified.

All isolates demonstrated effective degradation of glucose and utilized partially hydrolyzed laminarin more efficiently than HMW laminarin. Among the isolates, FI018 and FI145 showed the highest and lowest performance in substrate degradation, respectively.

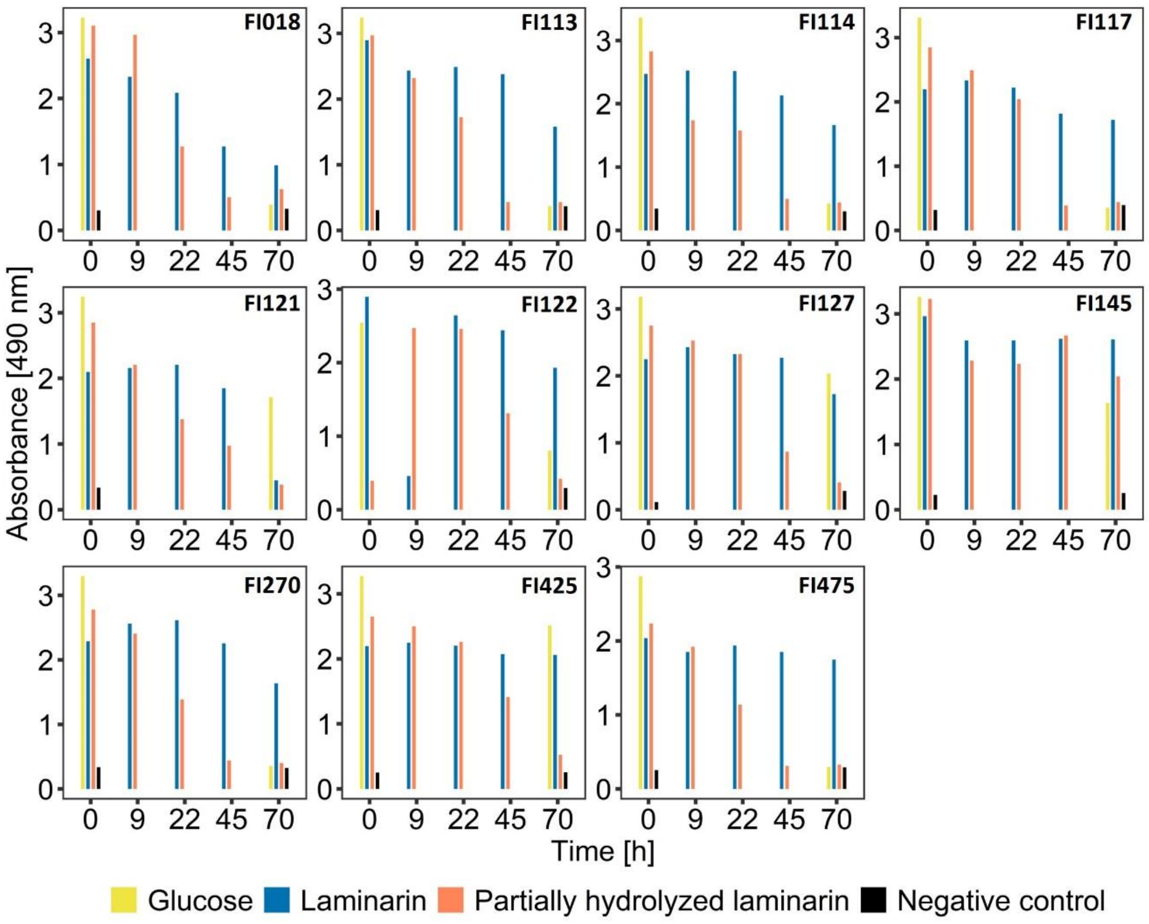
